# GRASP55 regulates mutant huntingtin unconventional secretion and aggregation

**DOI:** 10.1101/2021.10.05.463252

**Authors:** Erpan Ahat, Sarah Bui, Jianchao Zhang, Felipe da Veiga Leprevost, Lisa Sharkey, Whitney Reid, Alexey I. Nesvizhskii, Henry L. Paulson, Yanzhuang Wang

## Abstract

Recent studies demonstrated that the Golgi stacking proteins, GRASPs, especially GRASP55, regulate Golgi-independent unconventional secretion, but the underlying mechanism remains unknown. Here, we used mutant huntingtin (Htt-Q74) as a model system to address this question. Our results demonstrate that Htt secretion is GRASP55- and autophagy-dependent, and is enhanced under stress conditions such as starvation and ER stress. Mechanistically, GRASP55 facilitates Htt secretion by tethering autophagosomes to lysosomes to promote autophagosome maturation and by stabilizing p23/TMED10, a channel for translocation of cytoplasmic proteins into the ERGIC lumen. Moreover, GRASP55 level is upregulated by various stresses to facilitate unconventional secretion, while inhibition of Htt-Q74 secretion by GRASP55 knockout enhances Htt aggregation and toxicity. Lastly, comprehensive secretomic analysis identified novel cargoes secreted by the same unconventional pathway, such as TAGLN, PAICS and PRDX1. This study provides important information on the role of GRASP55 in unconventional protein secretion and Huntington’s disease progression.

## Introduction

Neurodegenerative diseases are characterized by increased neuronal cell death and decreased cognitive abilities of patients. Different diseases, such as Alzheimer’s disease (AD), Parkinson’s disease (PD), Huntington’s disease (HD) and Amyotrophic lateral sclerosis (ALS), are caused by different protein-based aberrations that affect different pathways; yet all cause neuronal cell death by forming inclusion bodies (Lee et al., 2005, Umeda et al., 2014, Arrasate and Finkbeiner, 2012). These include protein aggregates formed by hyperphosphorylated tau in AD (Umeda et al., 2014), mutant superoxide dismutase 1 (SOD1) (Dangoumau et al., 2014) and TAR DNA binding protein TDP43 in ALS (Johnson et al., 2009), and mutant huntingtin (Htt) in HD (Bates et al., 2015, Saudou and Humbert, 2016). Many of these neurotoxic proteins have been detected in the cerebrospinal fluid (CSF) of disease patients, indicating that these proteins could be secreted from cells (Wild et al., 2015, Mecocci et al., 1998, Winer et al., 2013). How the secretion affects protein aggregation and toxicity in neurons, and whether secretion facilitates the spreading of toxic proteins to neighboring cells, remain elusive.

In the conventional secretory pathway, transmembrane and secretory proteins, such as the amyloid precursor protein (APP), are synthesized by the endoplasmic reticulum (ER) and transported to the Golgi, from where they are targeted to their final destinations. However, many neurotoxic proteins, such as Htt, SOD1, α-synuclein, tau, and TDP-43, are synthesized as cytosolic proteins. Secretion of these proteins is independent of the traditional secretory pathway in which the Golgi functions as a trafficking center. For example, mutant Htt exists in two forms, an intracellular form in patient neurons, and an extracellular form in CSF (Wild et al., 2015) exported through a not well characterized secretory pathway (Trajkovic et al., 2017, Caron et al., 2021).

Little is known about the machinery that controls unconventional secretion, and the interplay between Golgi-dependent conventional trafficking and Golgi-independent unconventional secretion has not been explored. Although the unconventional pathway is independent of the Golgi, unconventional secretion of numerous cytosolic proteins reported so far requires the Golgi reassembly stacking protein of 55 kDa (GRASP55) (Lee et al., 2016, Rabouille and Linstedt, 2016, Villeneuve et al., 2018). GRASP55, and its homolog GRASP65, were first characterized as key Golgi stacking proteins (Barr et al., 1997, Wang et al., 2003). They both form *trans*-oligomers to link the Golgi cisternae into the unique stacked architecture (Xiang and Wang, 2010). Knockdown (KD) of GRASP55 and GRASP65 by RNAi, or knockout (KO) of both GRASPs by CRISPR/cas9, results in Golgi fragmentation (Xiang and Wang, 2010, Bekier et al., 2017). GRASPs were first linked to unconventional secretion of Acyl-CoA binding protein (AcbA/Acb1) in *Dictyostelium discoideum* and *Saccharomyces cerevisiae (Duran et al., 2010, Kinseth et al., 2007)*. In *Drosophila*, the sole GRASP protein is involved in unconventional trafficking of α-integrin at specific stages of fly development (Schotman et al., 2008). In mammalian cells, GRASPs are required for unconventional trafficking of Cystic Fibrosis Transmembrane Conductance Regulator (CFTR) and the cytokine IL-1β (Gee et al., 2011, Dupont et al., 2011). How GRASP55 regulates autophagosome-dependent unconventional secretion, and whether GRASP55 is required for secretion of neurotoxic proteins such as Htt, are unknown.

In this study, we report that mutant Htt is unconventionally secreted in a GRASP55-dependent manner. Htt secretion is elevated under stress conditions, including energy deprivation and nutrient starvation, inhibition of proteasomal degradation, and induction of ER stress, which also upregulate GRASP55 expression. GRASP55 facilitates Htt secretion through two actions, one is to accelerate autophagic flux by facilitating autophagosome and lysosome fusion, the other is to stabilize p23/TMED10, a protein channel in the ER-Golgi-intermediate compartment (ERGIC) that translocates cytosolic proteins into the ERGIC lumen for unconventional secretion. Lastly, we performed systematic proteomic and secretomic analysis of wild-type (WT) and GRASP55 KO (55KO) cells and identified a list of cytoplasmic proteins whose secretion depends on GRASP55.

## Results

### GRASP55 is required for unconventional secretion of mutant Htt

To determine if GRASP55 is required for the secretion of cytosolic proteins related to neurodegeneration, we expressed a number of neurotoxic proteins, including Htt exon 1 fragment with a long polyglutamine (Q74) stretch (GFP-Htt-Q74), WT SOD1 (GFP-SOD1), TDP43 (Flag-TDP43), and Tau (Flag-Tau), in WT and 55KO HeLa cells established earlier (Bekier et al., 2017). For Htt, we chose the Htt exon 1 fragment because it has been shown that this fragment is harder to degrade, more toxic than full-length Htt, and detected in the CSF with an elevated concentration in HD (Sathasivam et al., 2013, Chen and Wolynes, 2017, Fodale et al., 2017, Gerson et al., 2020). At 48 h post transfection, we cultured the cells in a serum-free medium for 4 h and determined the level of the corresponding cargo in the conditioned media by western blot. A conventional secretory pathway cargo, clusterin, was used as a control. All these proteins were found in the conditioned media (Figure S1A-D), but Htt-Q74 secretion was the most robust and GRASP55 dependent (Figure S1A). Therefore, we used GFP-Htt-Q74 as a main marker to study the mechanism of GRASP55-dependent unconventional secretion.

Since both GRASP55 and GRASP65 have been shown to be involved in unconventional secretion (Kim et al., 2016), we expressed GFP-Htt-Q74 in WT, 55KO, 65KO or DKO (double KO of both GRASP55 and GRASP65) HeLa cells. In this experiment GFP-Htt-Q74 expression did not cause cell death. At 48 h post transfection, about 90-95% cells contained soluble Htt in the cytosol, as indicated by the diffused GFP signal. Another 5-10% of cells contained Htt aggregates that appeared as bright dots in the cytoplasm (Figure 1A). Interestingly, these Htt aggregates were often found in the perinuclear region where the Golgi was localized. This prompted us to look at the effect of Htt expression on the Golgi morphology. While soluble Htt did not seem to affect Golgi morphology, Htt aggregates induced Golgi defects as previously reported (Sbodio et al., 2018, Brandstaetter et al., 2014), and the effect was more robust in 55KO cells (Figure 1A). For comparison, we also tested GFP-Htt-Q23 in parallel, which did not form aggregates or significantly affect the Golgi morphology (Figure S1E). Similarly, expression of SOD1, TDP43 or Tau in WT cells had no significant impact on the Golgi structure (Figure S1F).

**Figure 1.**
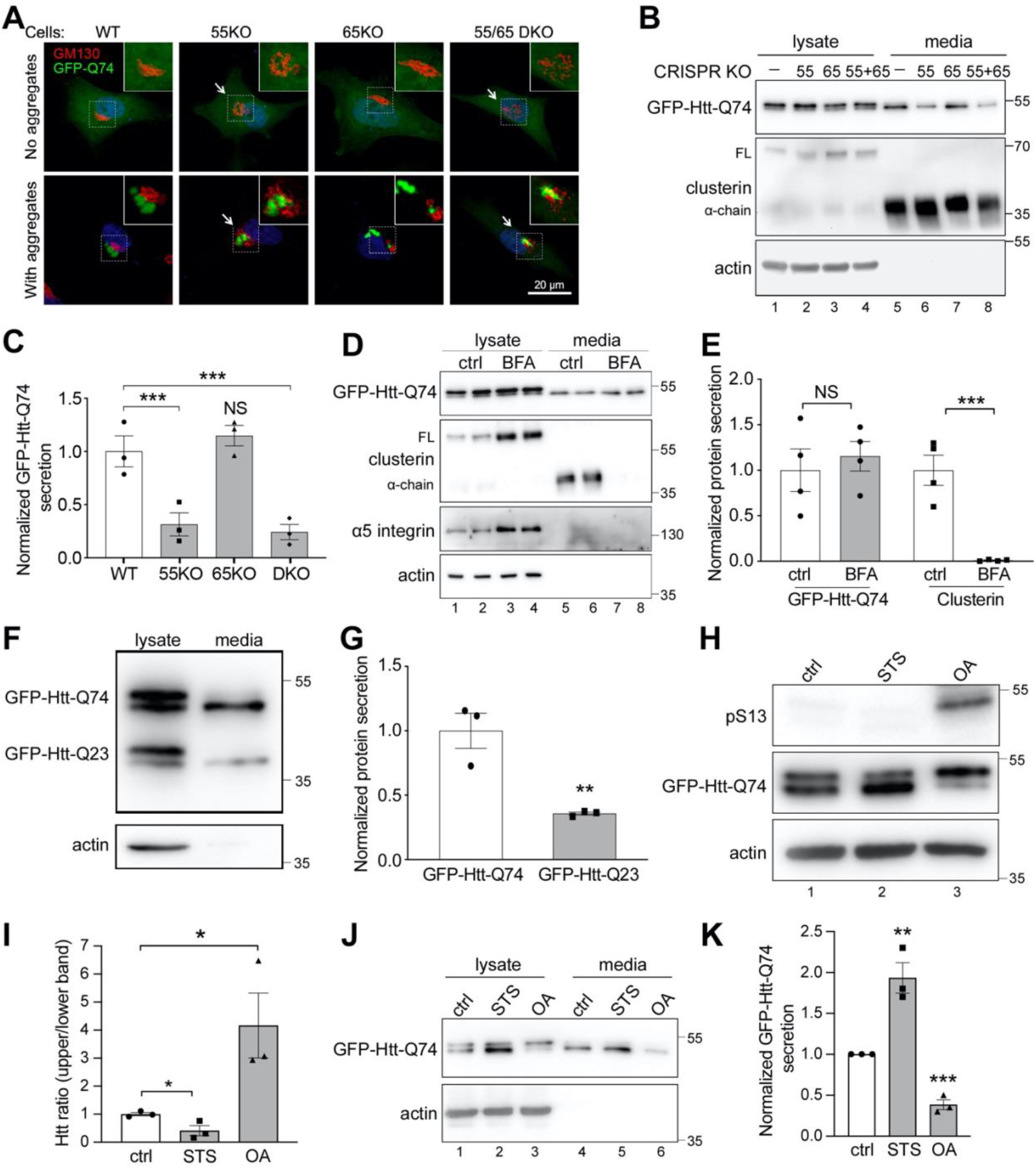
GRASP55 is required for unconventional secretion of mutant huntingtin. **(A)** GFP-Htt-Q74 aggregation induces Golgi fragmentation in GRASP55 KO (55KO) cells. WT and indicated GRASP KO HeLa cells expressing GFP-Htt-Q74 were stained for GM130 (red) and DNA (blue). Shown are example cells without (upper panels) or with (lower panels) Htt aggregates (arrows). **(B)** Htt secretion requires GRASP55 but not GRASP65. Cell lysates and conditioned media of GFP-Htt-Q74 expressing WT and GRASP KO cells were analyzed by western blot for Htt. Clusterin was used as a marker for conventional secretion. FL, full length clusterin; α-chain, mature clusterin. **(C)** Quantitation of Htt secretion in B. Htt secretion in WT was normalized to 1. **(D)** Htt secretion is unaffected by the block of ER-to-Golgi trafficking. GFP-Htt-Q74 expressing WT cells were treated with or without 5 μg/ml BFA for 4 h and Htt in the cell lysates and conditioned media were analyzed by western blot. Note that BFA treatment reduced the level of clusterin but not Htt in the conditioned media. **(E)** Quantitation of Htt and clusterin secretion in D, with their levels in control (ctrl) normalized to 1. **(F)** Htt-Q74 is preferably secreted than Htt-Q23. WT cells were cotransfected by GFP-Htt-Q23 and GFP-Htt-Q74 for 24 h. Cells were then incubated in serum-free medium for 4 h followed by western blot of Htt in the cell lysate and conditioned media. **(G)** Quantification of Htt-Q74 and -Q23 in the conditioned media, with Htt-Q74 secretion normalized to 1. **(H)** Htt is phosphorylated. GFP-Htt-Q74 expressing cells were treated with 1 µM staurosporine (STS) or 1 µM okadaic acid (OA) for 4 h and analyzed by western blotting for GFP and pS13-Htt. The pS13 antibody recognizes Htt phosphorylated at serine-13. **(I)** Quantification of Htt in H based on the GFP blot. **(J)** Dephosphorylated Htt is preferably secreted. Cells treated as in H were analyzed for GFP-Htt-Q74 secretion. **(K)** Quantification of J. Quantitation results are presented as mean ± SEM; statistical analysis was performed using Student’s *t*-test. *, p < 0.05; **, p<0.01; ***, p<0.001.

Next, we tested whether Htt secretion is GRASP dependent. After transfection of GFP-Htt-Q74 in WT, 55KO, 65KO and DKO cells for 24 h, we cultured the cells in a serum-free medium for 4 h and determined the level of Htt in the conditioned media by western blot. Htt was readily detectable in the conditioned media, and the level was significantly reduced in 55KO but not 65KO cells (Figure 1B-C), indicating that Htt secretion depends on GRASP55 but not GRASP65. In this assay, actin was not detected in the conditioned media, indicating that Htt was secreted rather than non-selectively released from dead cells, while the secretion and processing of clusterin, a cargo molecule secreted by the conventional secretory pathway, were not affected by GRASP KO. Taken together, GFP-Htt-Q74 expression causes Golgi defects especially in 55KO cells, while 55KO reduces GFP-Htt-Q74 secretion.

### Mutant Htt is selectively secreted through a Golgi-independent pathway

To confirm that Htt is secreted through an unconventional Golgi-independent pathway, we blocked ER-to-Golgi trafficking by brefeldin A (BFA) treatment. As shown in Figure 1D-E, BFA treatment diminished clusterin in the conditioned media, indicating a block of conventional secretion. Htt secretion was unaffected by BFA treatment, confirming that it is secreted via a Golgi-independent unconventional secretory pathway.

The toxicity of Htt is associated with the length of the polyQ track within the exon 1 fragment; Htt with more than ∼35 glutamines in the stretch is regarded as a disease mutant (Bates et al., 2015, Saudou and Humbert, 2016). To determine whether the length of the ployQ track affects Htt secretion, we co-expressed GFP-Htt-Q74 and GFP-Htt-Q23 in the same cells and compared their secretion. As shown in Figure 1F, when GFP-Htt-Q74 and GFP-Htt-Q23 were co-expressed, the level of GFP-Htt-Q74 in the conditioned media was higher than that of GFP-Htt-Q23.

Interestingly, Htt exhibited two bands, and only the lower band was detected in the conditioned media; we were curious about the difference between the two bands. Htt is highly modified by post-translational modifications such as phosphorylation, acetylation, and cleavage (Arbez et al., 2017). Among these, phosphorylation at S13 and S16 by TANK-Binding Kinase 1 (TBK1) or IκB kinase (IKK) has been shown to reduce Htt aggregation and toxicity (Thompson et al., 2009, Hegde et al., 2020). To determine whether the two bands represent different phosphorylation forms of Htt, we treated cells with a general serine/threonine kinase inhibitor, staurosporine (STS), or a general phosphatase inhibitor, okadaic acid (OA). STS treatment increased the intensity of the lower band, while OA treatment accumulated the upper band, suggesting that the upper band represents a phosphorylated form while the lower band is dephosphorylated. This conclusion was confirmed using a phospho-specific antibody pS13 for Htt (Hegde et al., 2020), which recognized only the upper band (Figure 1H-I). More importantly, STS treatment increased Htt secretion, whereas OA treatment reduced its secretion (Figure 1J-K), indicating that the dephosphorylated, aggregate-prone form of Htt is preferentially secreted compared to the phosphorylated, soluble form of Htt. Taken together, these results demonstrate that unconventional secretion is selective, at least to a certain degree, and may serve as a clearing mechanism for cells to expel toxic proteins.

### GFP-Htt-Q74 secretion is enhanced by cellular stress

It has been demonstrated that unconventional secretion is elevated under different stress conditions when the autophagy level is high (Giuliani et al., 2011), and that unconventional secretion of CFTR and Acb1 depends on autophagy or autophagy genes (Noh et al., 2018, Bruns et al., 2011). In addition, we discovered that GRASP55 functions as an energy sensor through O-GlcNAcylation to regulate autophagosome maturation. Under growth conditions, GRASP55 is O-GlcNAcylated and concentrated on the Golgi. Upon glucose starvation, GRASP55 is de-O-GlcNAcylated and relocated from the Golgi to the interface between autophagosomes and lysosomes, where it interacts with LC3 on autophagosomes and LAMP2 on lysosomes and functions as a membrane tether to facilitate autophagosome-lysosome fusion (Zhang et al., 2018). Similar responses of GRASP55 occur under amino acid starvation (Zhang et al., 2019). In addition, GRASP55 also interacts with Beclin-1 to facilitate the assembly and membrane association of the UVRAG phosphatidylinositol 3-kinase (PtdIns3K) complex, and thereby facilitates autophagosome maturation (Zhang et al., 2019).

These observations prompted us to test the effect of energy deprivation and nutrient starvation on Htt secretion. As expected, both glucose starvation (GS) and amino acid deprivation [E, by incubating cells in Earle′s Balanced Salt Solution (EBSS)] increased the level of Htt in the conditioned media compared to DMEM (D) (Figure 2A-B). A similar effect was observed in 55KO cells, although the overall secretion was significantly lower than WT cells. In addition to starvation, inhibiting proteasomal degradation with MG132, or inducing ER stress with tunicamycin (Tu), both enhanced Htt secretion (Figure 2C-F). These results confirm that Htt secretion is elevated under stress conditions.

**Figure 2.**
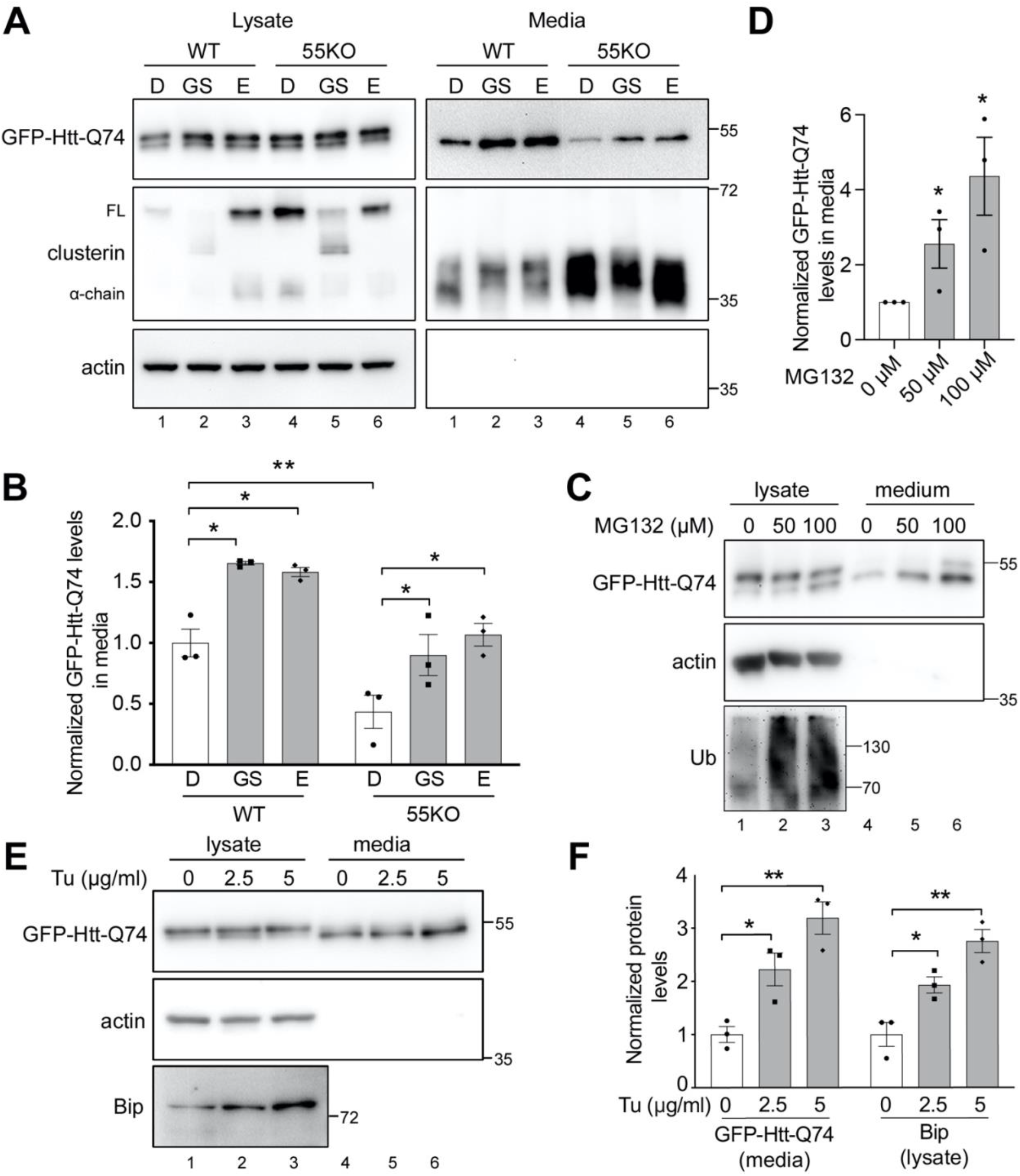
Htt secretion is enhanced by various stress stimuli. **(A)** Htt secretion is elevated by energy deprivation and nutrient starvation. WT and 55KO HeLa cells transfected with GFP-Htt-Q74 were incubated in serum-free DMEM (D), glucose free medium (GS) or EBSS (E) for 4 h followed by western blot of Htt and clusterin in the cell lysates and conditioned media. **(B)** Quantification of Htt secretion in A. (**C**) Htt secretion is enhances by proteostasis stress. WT HeLa cells expressing GFP-Htt-Q74 were treated with indicated concentration of MG132 for 4 h followed by western blot of Htt in the cell lysates and conditioned media. Ubiquitin (Ub) was used as a control for MG132 treatment. **(D)** Quantification of Htt secretion in C. **(E)** Htt secretion is enhances by ER stress. WT HeLa cells expressing GFP-Htt-Q74 were treated with indicated concentration of tunicamycin (Tu) for 4 h and analyzed by western blot for Htt. Bip was used as a marker for ER stress response. **(F)** Quantification of E. Results are presented as mean ± SEM; statistical analysis was performed using Student’s *t*-test. *, p < 0.05; **, p < 0.01.

### GRASP55 level is upregulated by cellular stress and by Htt expression

The fact that GRASP55 localization and function are regulated by cellular stressors suggests that GRASP55 may serve as both a stress sensor and effector (Zhang and Wang, 2020). In addition to its relocalization from the Golgi to autophagosomes under starvation conditions (Zhang et al., 2018), we wondered whether GRASP55 level might also be upregulated to meet the high demand for its roles in Golgi structure formation, autophagy, and unconventional secretion in response to stress. To test this hypothesis, we incubated WT HeLa cells in EBSS for 0, 2, 4 and 8 h and measured GRASP55 level by western blot. Indeed, EBSS treatment increased GRASP55 expression, but had no effect on other Golgi proteins we tested, including GRASP65, Golgin-160, Golgin-97, and GCC88 (Figure 3A-B). The elevation of GRASP55 expression was not limited to amino acid starvation, as it also applied to ER stress. When cells were treated with ER stress inducers, including Tu, thapsigargin (Tg) and DTT, the level of GRASP55 but not that of other Golgi proteins increased (Figure 3C-D).

**Figure 3.**
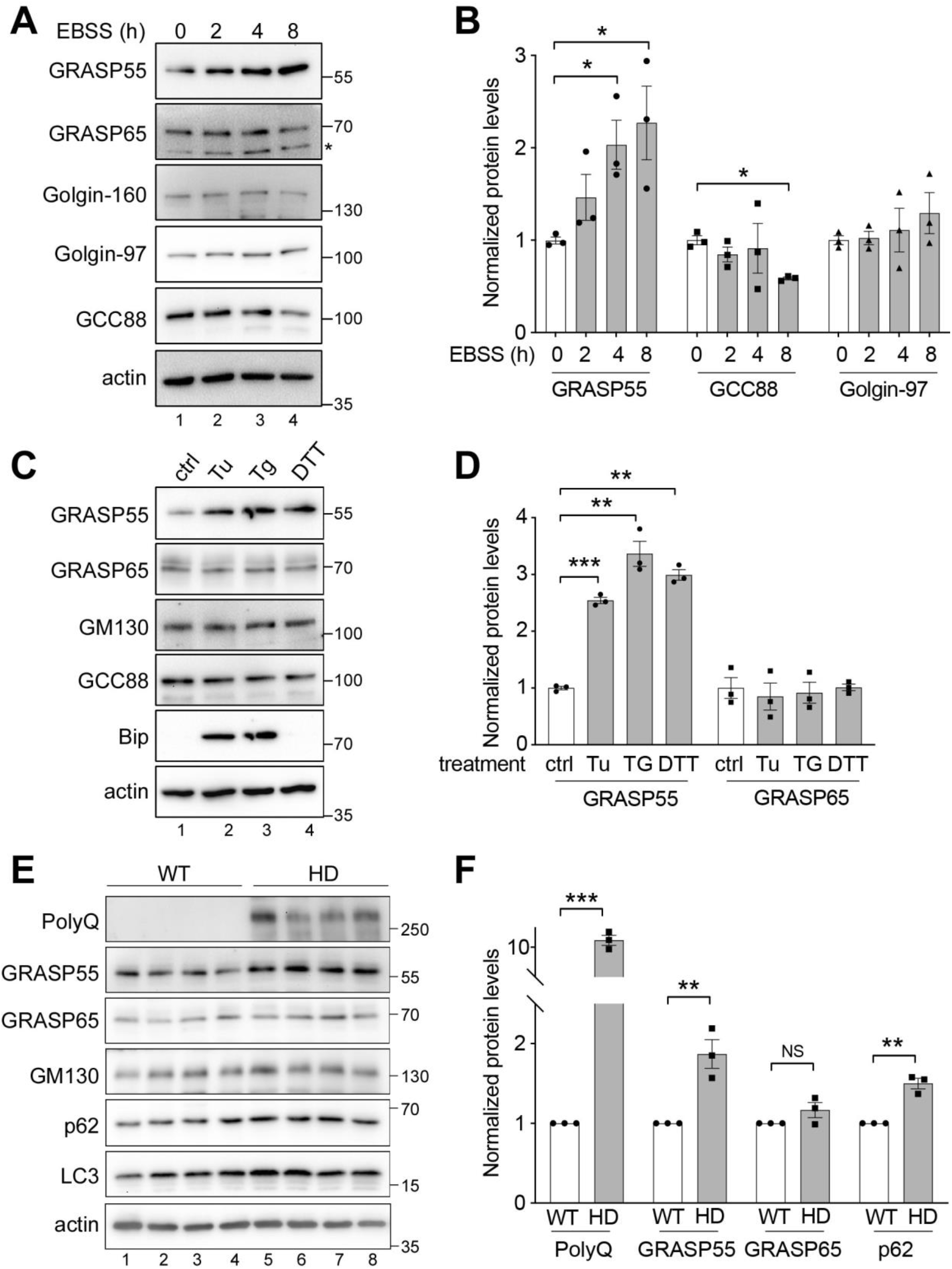
GRASP55 expression is upregulated under different stress conditions. **(A)** GRASP55 level is increased after amino acid starvation. WT HeLa cells were treated with EBSS for 0, 2, 4 or 8 h and analyzed for indicated Golgi proteins. Note that only GRASP55 is upregulated over time. *, nonspecific band. **(B)** Quantitation of indicated proteins in A. **(C)** GRASP55 expression is upregulated by ER stress. Cells were treated with 1 µg/µl tunicamycin (Tu), 1 µM thapsigargin (Tg) or 0.5 mM DTT for 24 h and probed for indicated Golgi proteins and an ER stress marker Bip. **(D)** Quantitation of GRASP55 and GRASP65 in C. **(E)** GRASP55 expression is elevated in Htt transgenic mice. Brain samples of 8-10 months old Htt transgenic mice (HD, Htt-Q200 expressed under the Htt protomer in knock-in mice) and WT littermates were analyzed by western blot of indicated proteins. Only GRASP55 increased its level in HD mice compared to WT. **(F)** Quantitation of indicated proteins in E. Results are presented as mean ± SEM; statistical analysis was performed using Student’s *t*-test. *, p < 0.05; **, p < 0.01; ***, p<0.001.

Since mutant Htt forms aggregates that could induce proteostasis stress, we tested the effect of Htt-Q23 and Htt-Q74 on GRASP55 expression. Both Htt-Q23 and Htt-Q74 expression increased the GRASP55 level in cells, although Htt-Q74 exhibited a stronger effect (Figure S1G-H). To validate this observation in mice, we took advantage of a previously established knock-in mouse model that expresses full-length Htt-Q200 in the brain (Heng et al., 2010, Gerson et al., 2020). These studies have shown that expression of Htt-Q200 upregulated both LC3-II and p62 levels compared to control, indicating an enhanced autophagosome formation but reduced lysosomal degradation. This conclusion was validated in our Htt mice and cell samples, as indicated by p62 accumulation (Figure 3E-F; Figure S1G-H) (Heng et al., 2010). In response to the proteostasis stress, GRASP55 level was also elevated in the brain tissues, while other proteins such as GRASP65 and GM130 did not change (Figure 3E-F). Taken together, GRASP55 level is upregulated by cellular stress and Htt expression, possibly to meet the high demand of unconventional secretion.

### GFP-Htt-Q74 secretion is autophagy dependent

The fact that GRASP55 expression, autophagy and unconventional secretion are all elevated under stress conditions implies that these events are related. To test if Htt secretion depends on autophagy, we expressed GFP-Htt-Q74 in the Atg7 knockout mouse embryonic fibroblast (MEF) cell line established by Dr. Masaaki Komatsu (Komatsu et al., 2005) in which autophagosome formation was abolished, as indicated by the lack of LC3-II (Figure 4A). Htt secretion was reduced in Atg7 KO cells to an almost undetectable level (Figure 4A-B). In addition, inhibition of autophagosome-lysosome fusion by bafilomycin A1 (BafA1) treatment also decreased Htt secretion in WT but not 55KO cells (Figure 4C-D).

**Figure 4.**
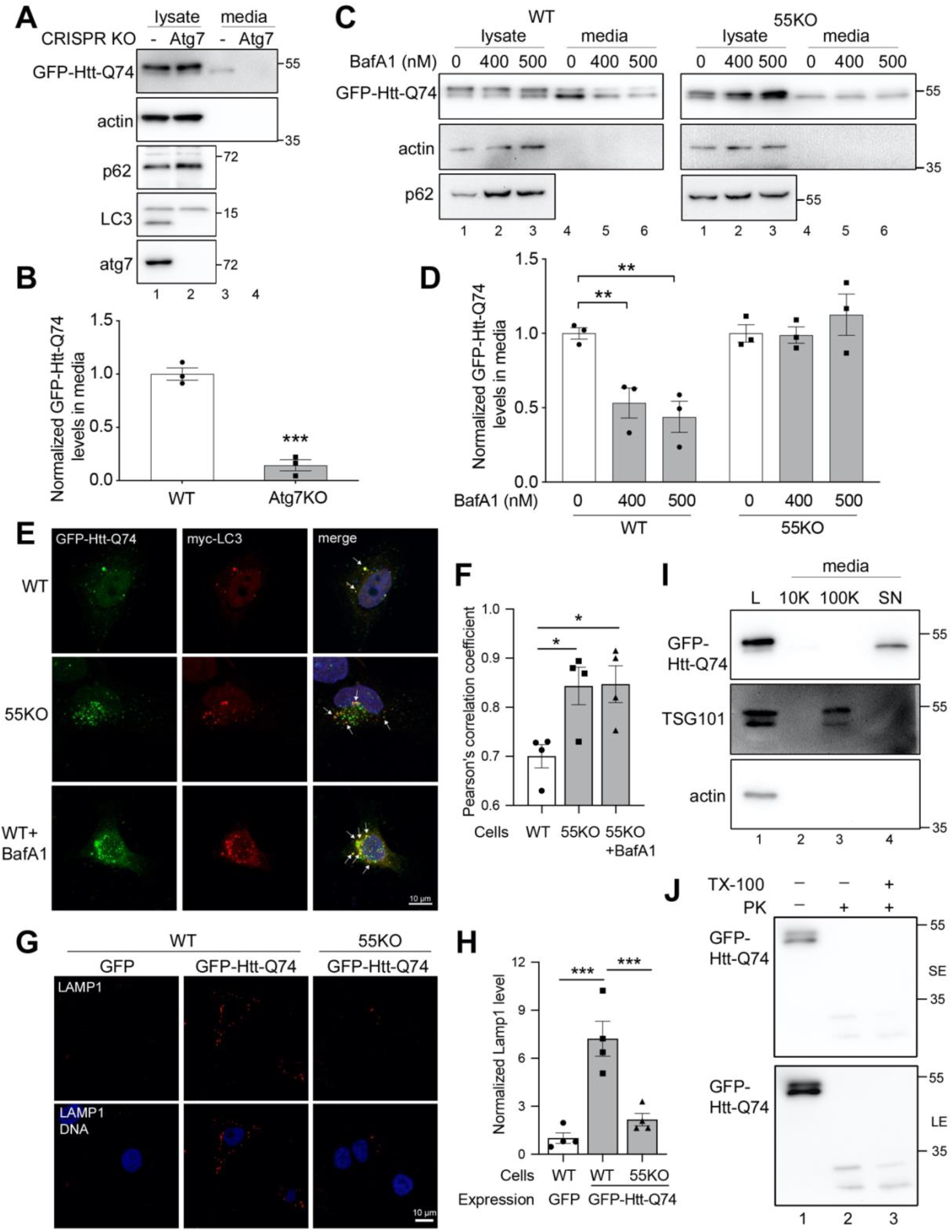
Htt secretion is autophagy dependent. **(A)** Htt secretion is inhibited by ATG7 KO. WT and ATG7 KO MEF cells were transfected with GFP-Htt-Q74 and analyzed for Htt secretion by western blot. **(B)** Quantitation of Htt secretion in A. **(C)** Htt secretion is reduced by autophagy inhibition. WT and 55KO HeLa cells expressing GFP-Htt-Q74 were treated with 0, 400, or 500 nM BafA1 for 4 h and analyzed for Htt secretion by western blot. Note that BafA1 reduced Htt secretion in WT but not 55KO cells. **(D)** Quantitation of Htt secretion in C. **(E)** Htt colocalizes with autophagosomes. Htt expressing cells were pre-permeabilized with 0.1% saponin for 2 min, fixed and immunostained with a myc antibody. **(F)** Quantitation of Htt and LC3 colocalization in E. **(G)** Htt is likely released by lysosome exocytosis. WT HeLa cells expressing GFP or GFP-Htt-Q74 were stained for LAMP1 without permeabilization. **(H)** Quantitation of cell surface LAMP1 level in G, with LAMP1 GFP-expressing WT cells normalized to 1. Results are presented as mean ± SEM; statistical analysis was performed using Student’s *t*-test. *, p < 0.05; **, p < 0.01; ***, p<0.001. **(I)** Htt secretion is independent of exosomes. Conditioned medium from GFP-Htt-Q74 expressing cells was harvested and subjected to differential centrifugation. Equal proportions of each pellet and soluble fraction were analyzed by western blot for actin, TSG101, and GFP. L, whole cell lysate; 10K, 10,000 *g* 30 min pellet; 100K, 100,000 *g* 60 min pellet; SN, 100,000 *g* 60 min supernatant. **(J)** Secreted Htt is not enclosed in membrane structures. Conditioned medium from GFP-Htt-Q74 expressing cells was treated with 2.5 μg/ml proteinase K (PK) for 10 min in the presence or absence of Triton X-100 followed by western blot of GFP. SE, short exposure; LE, long exposure.

The results described above are consistent with the role of GRASP55 in autophagosome-lysosome fusion, suggesting that GRASP55 regulates Htt unconventional secretion via controlling autophagosome maturation. If this conclusion is correct, we would expect to see more colocalization of Htt with the autophagosome marker LC3 in 55KO and BafA1-treated cells. Consistent with this idea, Htt partially colocalized with LC3 in WT cells, while 55KO and BafA1 treatment enhanced the colocalization (Figure 4E-F). It has been indicated that unconventional secretion could occur through lysosomal secretion (Villeneuve et al., 2018). Indeed, GFP-Htt-Q74 expression, but not GFP expression, enhanced the signal of the lysosome protein LAMP1 at the cell surface (Figure 4F), while this effect was highly reduced in 55KO cells (Figure 4G-H). These results indicate that Htt is probably taken up by autophagosomes, and once autophagosomes are fused with lysosomes, Htt can be released by lysosome exocytosis.

It has previously been shown that Htt is secreted in exosomes (Trajkovic et al., 2017), which is in conflict with our conclusion that Htt secretion is autophagy-dependent. Therefore, we performed differential centrifugation to pellet exosomes in the conditioned media and determined the level of Htt in different fractions. As shown in Figure 4I, exosomes were highly enriched in the pellet of the 100,000 *g* spin as indicated by the exosome marker TSG101 (lane 3), while Htt was only found in the supernatant (lane 4). To confirm that Htt was not secreted in membrane-bound structures such as exosomes, we treated the conditioned media of Htt expressing cells with proteinase K with or without Triton X-100 (TX-100). The results showed that Htt was degraded by proteinase K regardless of whether TX-100 was included (Figure 4J). Taken together, these results demonstrate that Htt secretion depends on autophagy but not exosome secretion.

### GRASP55 facilitates GFP-Htt-Q74 secretion through promoting autophagosome maturation and stabilizing p23/TMED10

GRASP55 facilitates autophagosome maturation by tethering autophagosomes and lysosomes through the interactions with LC3-II on autophagosomes and LAMP2B on lysosomes (Zhang et al., 2018). To confirm that GRASP55 promotes Htt secretion through regulating autophagosome maturation, we re-expressed GRASP55 and its autophagy mutants in 55KO cells and tested their effect on Htt secretion. Expression of WT GRASP55 restored GFP-Htt-Q74 secretion, showing that the reduced Htt secretion in 55KO cells was specific for GRASP55 depletion (Figure 5A). In contrast, expression of the F37A mutant of GRASP55, which abolishes GRASP55-LC3 interaction (Zhang et al., 2018), failed to rescue the secretion defect of Htt in 55KO cells (Figure 5B-C). The 5A mutant of GRASP55, whose O-GlcNAcylation is disabled and thus its role in autophagy is constitutively active, also rescued Htt secretion similar to WT GRASP55. In this experiment, GRASP55 itself is not secreted with Htt; and we did not detect an interaction between Htt and GRASP55 by co-immunoprecipitation. These results demonstrate that GRASP55 facilitates Htt secretion by promoting autophagosome maturation.

**Figure 5.**
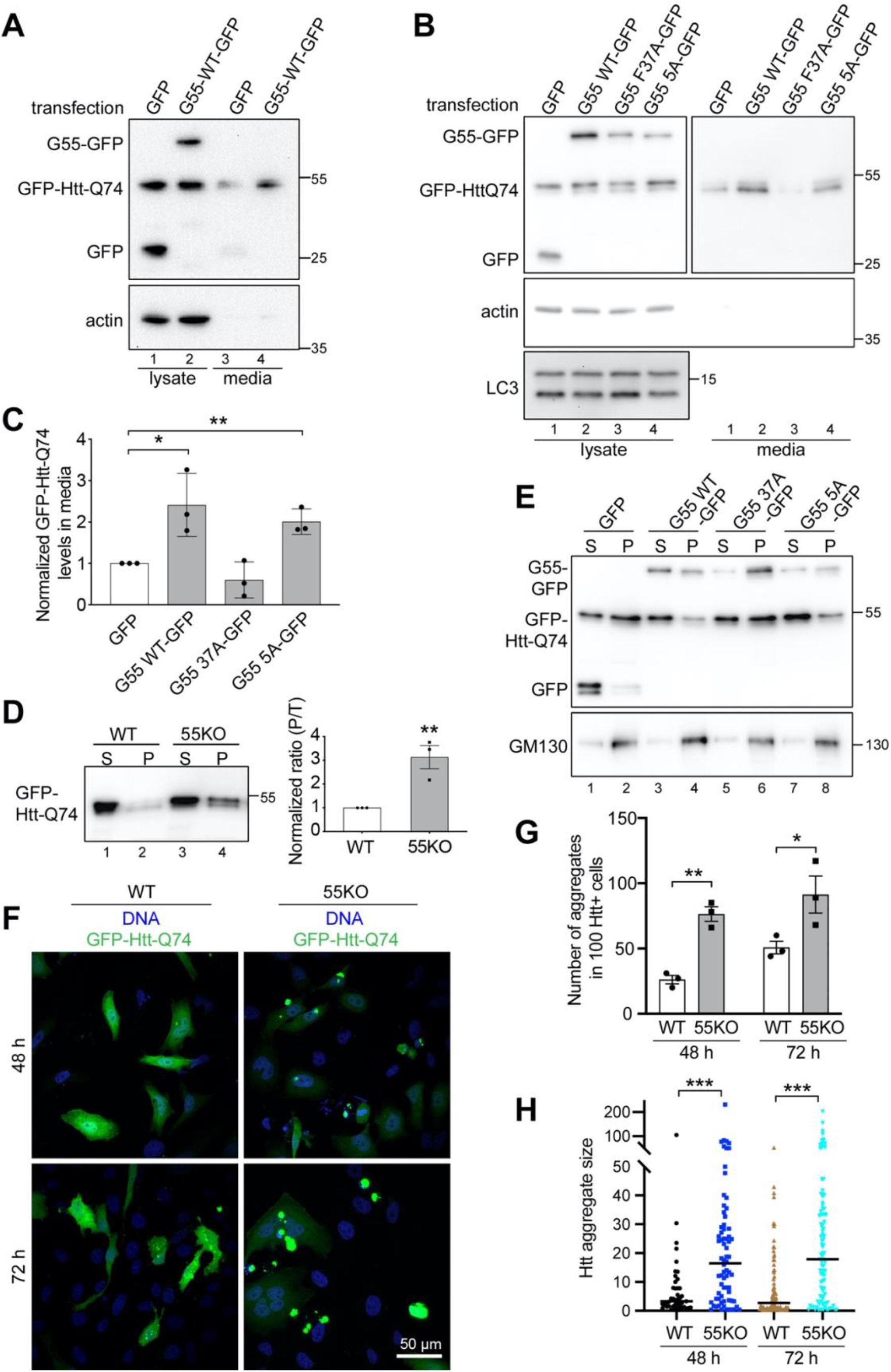
GRASP55 regulates Htt secretion via autophagy and p23. **(A)** GRASP55 expression rescues the defect of Htt secretion in 55KO cells. 55KO cells were first transfected with GFP-Htt-Q74 for 24 h, and then with either GFP or WT GRASP55-GFP for another 24 h. Htt secretion was analyzed by western blot. **(B)** Expression of WT GRASP55 but not its autophagy inactive F37A mutant rescues the defect of Htt secretion in 55KO cells. GFP-Htt-Q74 expressing 55KO cells were transfected with GFP, WT GRASP55-GFP or its F37A or 5A mutant for 24 h. Htt secretion was analyzed by western blot. **(C)** Quantitation of Htt secretion in B. **(D)** Htt aggregation is enhanced by 55KO. WT and 55KO cells were transfected by GFP-Htt-Q74 for 48 h and lysed by 0.5% NP-40 followed by centrifugation at 136,000 *g* at 4ºC for 1 h. Supernatant (S) and pellet (P) were loaded at a 1:6 ratio and blotted for GFP. T, total. **(E)** Expression of WT GRASP55 but not F37A reduces Htt aggregation in 55KO cells. 55KO cells transfected as in B were analyzed for Htt aggregation as in D. **(F)** Htt forms more aggregates in 55KO than WT cells. WT and 55KO cells were transfected with GFP-Htt-Q74 for 48 h (upper panels) or 72 h (lower panels), stained for DNA, and imaged. **(G)** Quantification of the number of Htt aggregates in F. (**H)** Quantification of the size of Htt aggregates in F. Htt aggregate size quantification is performed using NIS-Elements software simple ROI editor function. Results are presented as mean ± SEM; statistical analysis was performed using Student’s *t*-test. *, p < 0.05; **, p < 0.01; ***, p<0.001.

Recently it has been reported that an ERGIC protein p23, also known as TMED10, functions as a protein channel for cytosolic proteins to be translocated into the ERGIC lumen before they are transported to autophagosomes and secreted to the extracellular space (Zhang et al., 2020). Indeed, p23 depletion largely reduced GFP-Htt-Q74 secretion (Figure S2A). Considering that some members of the p24 family proteins interact with GRASP65 (Barr et al., 2001) and that GRASP55 is partially relocated to the ER under stress conditions for CFTR trafficking (Kim et al., 2016), inhibition of Htt secretion by p23 depletion and 55KO suggested that GRASP55 may function together with p23 in Htt secretion. This speculation was first supported by the observation that the p23 level was reduced in 55KO but not 65KO cells (Figure S2B-C). In addition, p23 co-immunoprecipitated with GRASP55 but not GRASP65, while known interactions between GRASP65 and GM130 and between GRASP55 and Golgin-45 were readily detectable in the same experiment (Figure S2D). These results support the idea that GRASP55 interacts with p23 to facilitate Htt unconventional secretion.

### Inhibition of Htt secretion by 55KO enhances Htt aggregation and toxicity in the cell

It is generally believed that HD is caused by the accumulation of toxic mutant Htt protein in the cell. To determine the functional significance of Htt secretion, we assessed Htt aggregation in WT and 55KO cells. We lysed GFP-Htt-Q74 expressing cells in detergent and separated Htt aggregates from its soluble form by centrifugation. Western blot analysis of the two fractions showed that Htt formed more aggregates in 55KO than WT cells (Figure 5D). Expression of WT GRASP55 and its 5A mutant, but not the autophagy inactive F37A mutant, reduced Htt aggregation in 55KO cells (Figure 5E), consistent with their roles in Htt secretion. In this experiment, a significant amount of GRASP55 and GM130 were found in the pellet (Figure 5E), consistent with the idea that they form a detergent-insoluble Golgi matrix (Xiang and Wang, 2011, Wang and Seemann, 2011). We also confirmed the effect of GRASP55 KO on Htt aggregation by microscopy. At both 48 h and 72 h post-transfection, 55KO cells showed more and larger aggregates compared to WT (Figure 5F-H). These results demonstrate that GRASP55 dependent Htt secretion reduces its aggregation and toxicity in the cell.

### Secretome analysis of WT and 55KO cells identifies new proteins secreted by GRASP55-dependent unconventional secretion

To identify native proteins that are secreted in a GRASP55-dependent mechanism, we analyzed the conditioned media of WT and 55KO cells by Tandem Mass Tag (TMT) labeling and liquid-chromatography mass spectrometry (LC-MS). With a 1% false discovery rate (FDR), we identified 1696 proteins in the media (Table S1). We plotted all 1696 secreted proteins on a volcano plot to display their changes in the 55KO vs. WT secretome (Figure 6A). Significantly changed cargoes [|Log2FC (55KO/WT)|>0.5, p < 0.05] without predicted signal peptides are shown in black. On this plot, majority of the unconventional targets were reduced in 55KO.

**Figure 6.**
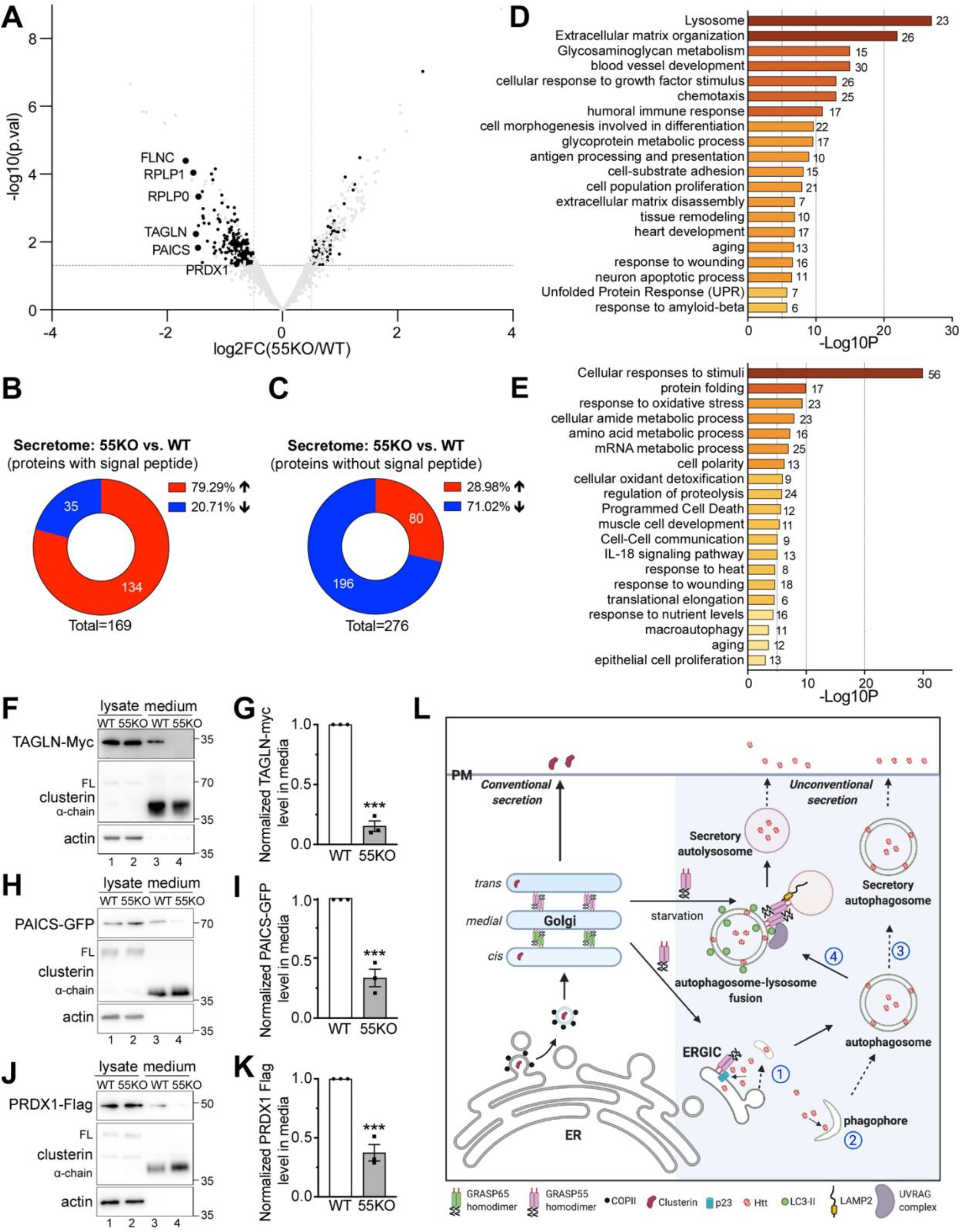
Secretome analysis identifies candidate proteins secreted by GRASP55-dependent unconventional secretion. **(A)** Volcano plot of candidate proteins in 55KO vs. WT secretome. Black dots represent significantly altered cargoes without signal peptide. All other cargoes are shown in gray. A few top candidates (large black dots) in the unconventional pathway are labeled. **(B)** GRASP55 KO increases overall conventional secretion. Indicated are the numbers of candidate proteins with ER signal sequences whose secretion was significantly increased or decreased in 55KO compared to WT cells [|Log2FC (55KO/WT)|>0.5; p < 0.05]. **(C)** GRASP55 KO decreases overall unconventional secretion. Number and percentage of candidates with no signal sequence that are increased or reduced in GRASP55 KO secretome compared to WT. **(D)** Metascape Gene Ontology (GO) enrichment analysis of identified candidate proteins with predicted ER signal sequences whose secretion was significantly altered in 55KO compared to WT. The number of proteins in each pathway is indicated on the right of the graph. **(E)** GO enrichment analysis of candidate proteins without predicted ER signal sequences whose secretion was significantly altered. **(F-K)** TAGLN, PAICS and PRDX1 are secreted in a GRASP55 dependent manner. Western blot and quantitation of each protein are shown. Results are presented as mean ± SEM; statistical analysis was performed using Student’s t test. ***, p<0.001. **(L)** GRASP55 functions in conventional and unconventional protein secretion. In the conventional secretory pathway (left), secretory proteins such as clusterin are transported from the ER to the Golgi and then to the plasma membrane (PM). GRASP55 depletion-mediated Golgi unstacking enhances the overall secretion of cargoes in the conventional pathway, as demonstrated in Figure 6C. In the unconventional pathway induced by stress (right), cytosolic proteins such as Htt may enter autophagosomes either via p23/TMED10-mediated translocation into the ERGIC lumen (1), from where it is incorporated into autophagosomes, or by direct sequestration into phagophores (2). Htt in autophagosomes can secreted directly by autophagosome exocytosis (3), or first transferred to lysosomes by membrane fusion (4) and released by lysosome exocytosis. Upon starvation, GRASP55 is relocated from the Golgi to 1) the autophagosome and lysosome interface to facilitates their fusion; 2) ERGIC to interact with p23/TEMD10 and regulates cytosolic protein translocation into the lumen. Our results that depletion of p23 and GRASP55 both reduce Htt secretion indicate that the dominant pathway for Htt translocation and secretion is 1 and 4.

Next, we systematically analyzed the impact of GRASP55 depletion on the secretion of both conventional and unconventional pathway cargoes. Among the 1696 proteins, we identified a total of 445 proteins whose secretion is highly affected by 55KO [|Log2FC (55KO/WT)|>0.5, p < 0.05] (Table S1). Within these 445 significantly changed proteins, 169 contained an ER signal sequence predicted by the SignalP-5.0 Server (http://www.cbs.dtu.dk/services/SignalP), while 276 proteins did not appear to have ER signal peptides (Table S1). Within the 169 proteins with ER signal sequences, 134 (79%) of them were increased in 55KO secretion (Figure 6B), consistent with our previous findings that GRASP depletion accelerates protein trafficking (Xiang et al., 2013). In contrast to the proteins with ER signal sequence, 196 (71%) of the 276 proteins without ER signal sequences were reduced in 55KO secretion (Figure 6C). It is worth mentioning that a GRASP55-dependent secretome has recently been reported (Nuchel et al., 2021). But unlike this report in which all proteins, including those with (e.g., MMP2) and without signal peptides, are included in unconventional secretion, we further differentiated the proteins into two separate groups and showed different effects by 55KO. Taken together, these results support our conclusion that GRASP55 plays essential roles in unconventional protein secretion.

To identify the pathways affected by 55KO, we performed Gene Ontology (GO) analysis of the significantly changed proteins. GO term pathway analysis of the 169 proteins with predicted ER signal peptides identified the following pathways as significantly changed in 55KO: lysosomal enzymes and structural proteins, extracellular matrix organization, glycosaminoglycan metabolism, cell adhesion and proliferation, cellular response to growth factor stimuli, immune response and tissue remodeling, and response to amyloid-beta, unfolded protein response and aging (Figure 6D; Table S2). The most affected pathway is related to lysosomal biogenesis, consistent with our previous finding that Golgi unstacking by GRASP depletion causes missorting and increased secretion of lysosomal enzymes (Xiang et al., 2013). Given the central role of the Golgi in protein trafficking, glycosylation, and secretion, it is also reasonable to see that GRASP55 depletion significantly alters extracellular matrix organization and glycosaminoglycan metabolism. In addition, our secretome analysis also validated our previous findings that 55KO affects cell adhesion, migration, and growth (Ahat et al., 2019).

GO term analysis of the 276 significantly changed proteins without signal sequences identified pathways such as cellular response to stimuli, oxidative stress, nutrient levels, heat, wounding, protein folding, macroautophagy, programmed cell death and cell-cell communication (Figure 6E; Table S2), which are consistent with the role of GRASP55 in stress response (Zhang and Wang, 2020). These findings further suggest a link between GRASP55, autophagy dependent-unconventional secretion and stress response. Consistent with this notion, previously identified unconventional secretion cargoes such as SOD1 and IL-β1 are related to oxidative stress and inflammation, and their secretion is increased under certain stress conditions such as nutrient starvation (Cruz-Garcia et al., 2017, Dupont et al., 2011).

To confirm that the identified proteins without ER signal sequences are indeed secreted through a GRASP55-depedent mechanism, we tested a few top candidates (Figure 6A), including transgelin 1 (TAGLN), multifunctional protein ADE2 (PAICS) and peroxiredoxin-1 (PRDX1), using the secretion assay we established for Htt. These proteins were selected because they are predicted to be highly secreted by the SecretomeP 2.0 server (http://www.cbs.dtu.dk/services/SecretomeP), an online tool to predict the probability of unconventional protein secretion, although their secretion and the GRASP55-dependence in the secretion have not been experimentally confirmed. Indeed, these proteins were all detected in the conditioned media, and their secretion was all reduced in GRASP55 KO cells (Figure 6F-K). In summary, the unbiased secretomic analysis provided new evidence that GRASP55 depletion accelerates conventional protein trafficking and secretion but inhibits unconventional secretion. It also identified a list of bona fide candidates that are secreted by GRASP55-dependent unconventional secretion.

## Discussion

In this study, we used GFP-Htt-Q74 as a model protein to study the role of GRASP55 in unconventional secretion and cytosolic aggregation of neurotoxic proteins. We discovered that mutant Htt is secreted through a GRASP55- and autophagy-dependent but Golgi-independent pathway. Our results also revealed a link between enhanced Htt secretion, elevated autophagy, and increased GRASP55 expression under various stress conditions, and confirmed a novel role of GRASP55 in stress response (Zhang and Wang, 2020). Htt may enter autophagosomes through p23-mediated translocation or by direct sequestration by phagophores. Due to lysosomal dysfunction, undegraded proteins in autophagosomes and lysosomes could be released by autophagosome or lysosome exocytosis (Figure 6L). GRASP55 enhances Htt secretion through two actions, one is to facilitate autophagosome-lysosome fusion by interacting with LC3 and LAMP2, and another is to stabilize p23, the translocon that allows cytosolic proteins enter the lumen of the ERGIC, from where they are transported to autophagosomes. Lastly, using systematic secretomic analysis, we identified and catalogued new proteins in GRASP55-dependent unconventional secretion.

It is interesting to see that GRASP55-dependent unconventional secretion is selective, at least to some degree. First, among all marker proteins tested, Htt and SOD1 are preferably secreted than TDP-43 and Tau. Consistently, the secretion of Htt and SOD1, but not TDP-43 and Tau, depends on GRASP55. It has been previously shown that Tau is secreted via direct translocation across the plasma membrane, which is regulated by heparan sulfate and *PtdIns(4,5)P*_*2*_ (Merezhko et al., 2018). So far there is no indication that GRASP55 is involved in this pathway. TDP43, on the other hand, has been shown to be secreted in a prion-like manner via exosomes (McAlary et al., 2020). While it is not clear whether GRASP55 directly regulates exosome secretion, our data indicates that it plays no role in TDP43 secretion. Second, Htt-Q74 is preferentially secreted than Htt-Q23. Given that Htt-Q74 forms more aggregates than Htt-Q23, one possible explanation is that aggregated proteins are selectively secreted. This speculation is supported by the third observation that of the two bands of Htt-Q74, only the aggregation-prone, dephosphorylated form is secreted. Whether Htt-Q74 aggregates are preferably enclosed by autophagosomes or selectively translocated into the ERGIC lumen requires further investigation.

In HD, mutant Htt with a long polyQ stretch tends to form toxic protein aggregates that cause cellular toxicity and neurodegeneration (Bates et al., 2015, Leitman et al., 2014, Arrasate and Finkbeiner, 2012). In this study we used the N-terminal fragment of Htt as a cargo due to its relevance to the disease. The exon 1 fragment of Htt contains the polyQ stretch which is the source of the toxicity of mutant Htt. This fragment is harder to degrade, more toxic than full-length Htt, and detected in the CSF with an elevated concentration in HD (Sathasivam et al., 2013, Chen and Wolynes, 2017). Like full-length Htt, the aggregation and toxicity of the exon 1 fragment are regulated by phosphorylation of S13 and S16, which indicates similar regulation. For these reasons we believe that the N-terminal fragment of Htt serves as an excellent reporter in our study.

Autophagy is an energy and nutrient deprivation-induced, lysosome-mediated degradation pathway essential for organelle turnover and degradation of aggregated proteins. Compared to non-neuronal cells, neurons need to maintain high basal autophagy for survival (Mizuno et al., 2001, Fujita et al., 2002). Autophagy and autophagy defects have been linked to neurodegenerative diseases (Hegde et al., 2020, Levine and Kroemer, 2008). While wildtype Htt is required for selective autophagy (Rui et al., 2015), mutant huntingtin has been shown to hinder autophagy flux, and certain population of toxic Htt is immune to autophagy-dependent degradation (Fu et al., 2017). It is possible that defects in lysosomal function and impairment of protein degradation may activate GRASP55-dependent unconventional secretion to clear out the toxic, autophagy-resistant Htt aggregates. Indeed, our data demonstrates that inhibition of proteasomal degradation pathway also significantly increase unconventional secretion of mutant Htt.

Htt secretion could be a double-edged sword to the neuronal cells and the disease. On the one hand, the aggregation-prone, dephosphorylated Htt with a long polyQ stretch is selectively secreted, which reduces the amount of Htt aggregates in the cells and subsequently ameliorates its toxicity in the cell. In this sense Htt secretion is beneficial to the cell. Our results support this hypothesis as GRASP55 depletion increases overall Htt aggregation in the cell (Figure 5D-H). On the other hand, secreted Htt aggregates may serve as seeds for internalization into the neighboring cells to trigger more Htt aggregation. Here Htt secretion may facilitate the spreading and propagation of Htt aggregates from disease to healthy neurons, which may contribute to the development of the disease. This hypothesis, however, is unlikely since we observed overall significantly more and larger Htt aggregates in GRASP55 KO cells compared to WT. Future studies will test the biological significance of Htt secretion through the intervention of GRASP55 in HD animal models. New information provided in this study may shed light on novel therapeutic approaches for the treatment of this devastating disease.

## Materials and Method

### Reagents, plasmids, and antibodies

All reagents were purchased from Sigma-Aldrich, Roche, Calbiochem, and Fisher unless otherwise stated. The following cDNA constructs have been described previously: pEGFP-N1-GRASP55 (WT, 37A, 5A) and pEGFP-N1-GRASP65 (WT) (Zhang et al., 2018). Others cDNAs include GFP-Htt-Q74 (Addgene, Cat# 40262), GFP-Htt-Q23 (Addgene, Cat # 40261), TAGLN-Myc (OriGene, Cat # RC215789), PAICS-EGFP (Addgene, Cat #99108), pFRT/TO/HIS/FLAG/HA-PRDX1 (Addgene, Cat # 38086).

Antibodies used in this study include monoclonal antibodies against β-actin (Proteintech, Cat# 66009-1-Ig), Clusterin (Santa Cruz, Cat# SC-5289), Flag (Sigma, Cat# M1804), GFP (Proteintech, Cat# 66002-1-lg), Myc (9E10, Dr. David Sheff) and PolyQ (Sigma, Cat# MAB1574); polyclonal antibodies against Atg7 (Cell Signaling, Cat# 2631S), Bip (Santa Cruz, Cat# sc-13968), GCC88 (Proteintech, Cat# 16271), Golgin-97 (Proteintech, Cat# 12640-1-AP), Golgin-160 (Proteintech, Cat#2 1193), GRASP55 (Proteintech, Cat# 10598-1-AP), human GRASP65 (Dr. Joachim Seemann, University of Texas Southwestern Medical Center), α5 integrin (Cell Signaling, Cat# 4705), LC3 (Cell Signaling, Cat#2755), and p62 (Proteintech, Cat# 18420-1-AP).

### Cell culture, transfection, and treatment

For cell culture, HeLa and MEF cells were grown in DMEM (Invitrogen) containing 10% super calf serum (Gemini, Cat# 100-510) and glutamax at 37°C in a 5% CO2 incubator. For Htt expression, GFP-Htt-Q74 was transfected into WT or GRASP KO HeLa cells in 3.5 cm dishes when the cells reached the confluency of 50-60%. For drug treatment, 5 µg/ml BFA, 500 ng/ml nocodazole, 100 µg/ml CHX, 50 µM MG132, 1 µM staurosporine, 1 µM okadaic acid or 5 µg/ml tunicamycin was directly applied to the medium for the indicated times.

### Protein secretion assay and TCA precipitation

WT or GRASP KO HeLa cells grown in 3.5 cm dishes were transfected with GFP-Htt-Q74 (4 µg DNA for a 3.5 cm dish) when the cells reached the confluency of 50-60% and incubated in full growth medium for 24 h. Cells were washed with phosphate buffered saline (PBS) five times and further incubated in 1 ml DMEM without serum for 4 h. Conditioned media were collected, cleared by centrifugation at 500 *g* for 10 min at 4ºC and 2000 *g* for 20 min at 4ºC. 0.9 ml of precleared conditioned medium was then mixed with 0.11 volumes of ice-cold 100% TCA (2.2 g/ml) and placed on ice. After 10 min, 500 μl of ice-cold 10% TCA were added to the sample-TCA mix. After 20 min samples were centrifuged at 20,000 *g* for 30 min at 4ºC. Pellet was washed with cold acetone (−20ºC), dried, and dissolved in 1x SDS loading buffer for western blot analysis. The cells were washed with cold PBS twice, scraped and lysed in 20 mM Tris-HCl, pH 8.0, 150 mM NaCl, 1% Triton X-100 and protease inhibitor cocktail for 30 min on ice. Lysates were cleared by centrifugation (20,000 *g* for 20 min at 4ºC). The cell lysate and medium precipitate were loaded at 1:8 ratio. After electrophoresis and transfer, nitrocellulose membranes were incubated with corresponding primary antibodies overnight at 4°C. The membranes were extensively washed and further incubated with HRP conjugated goat anti-rabbit or goat anti-mouse secondary antibodies for 1 h at room temperature and exposed to a FluorChem M machine (Proteinsimple).

### Immunofluorescence microscopy

Cells were grown on sterile glass coverslips and rinsed with PBS before fixation. To stain total cellular proteins, cells were fixed in 4% paraformaldehyde (PFA) for 10 min and permeabilized with 0.2% Triton X-100 in PBS for 10 min. Cells were incubated with primary antibodies overnight at 4ºC, washed and probed with the appropriate secondary antibodies conjugated to TRITC for 45 min. To stain cell surface LAMP1, cells were fixed with 1% PFA for 10 min without permeabilization. Cells were incubated with an anti-LAMP1 antibody (H4A3, DSHB, 1:100) overnight at 4ºC, washed and probed with the appropriate secondary antibodies conjugated to TRITC for 45 min at room temperature. To analyze LC3 and Htt colocalization, cells were pre-permeabilized in KHM buffer (125 mM potassium acetate, 25 mM Hepes, pH 7.2, and 2.5 mM magnesium acetate) containing 0.1% saponin (prefiltered) for 2 min on ice to release the cytosol and washed by KHM buffer for 5 min at room temperature. Cells were then fixed with 4% PFA, permeabilized in 0.2% Triton X-100 and incubated with primary and secondary antibodies as described above. DNA was stained with Hoechst in PBS for 5 min. Coverslips were rinsed with PBS and mounted with Mowiol onto slides. Images were taken with a 20x air objective or a 63x oil objective on a Nikon ECLIPSE Ti2 Confocal microscope and shown as max projections.

### Protease K protection assay

Protease K protection assay of the conditioned media was performed as previously described (Zhang et al., 2019). Briefly, conditioned medium from HeLa cells expressing GFP-Htt-Q74 was collected as described above and was equally divided into three tubes, one was left untreated on ice, the second was incubated with 2.5 μg/ml Protease K (Thermo Fisher Scientific, Cat # AM2542) on ice, and the third was treated with both protease K and 1% Triton X-100 (from 20% stock) for 10 min on ice. Protease K was then inhibited by adding 1 mM PMSF (from 100 mM stock in isopropanol) and further incubated on ice for 10 min. Proteins in each sample were pelleted by TCA precipitation as above. The pellets were dissolved in 1x SDS sample buffer (50 mM Tris-HCl, pH 6.8, and 2% SDS) and analyzed by western blotting.

### Separation of insoluble Htt aggregates from its soluble forms

WT or GRASP KO cells were transfected with GFP-HttQ74 for indicated times. Cells were washed three times with cold PBS, scraped in cold PBS, centrifuged at 1000 *g* for 3 min at 4ºC, and lysed in lysis buffer (20 mM Tris 8.0, 150 mM NaCl, 0.5% NP40, protease inhibitors) for 20 min on ice. The supernatant and pellet were collected after spinning down at 55,000 rpm for 1 h. The supernatants concentration was adjusted to 1.5 mg/ml (the volumes of the samples are different) and supplemented with 1/5 volume of 6x SDS loading buffer with DTT (final 1x SDS in the sample). The pellet was dissolved in the same amount of 2x SDS loading buffer with DTT. Both samples were boiled, and the pellet and supernatant were loaded onto the gel at a 6:1 ratio.

### Proteomics analysis

WT and 55KO cells were cultured in full growth medium in 15 cm dishes in triplicates till they reach 80% confluency. The medium was exchanged to 20 ml DMEM only (without serum) for 12 h, cells and conditioned media were collected. The media were cleaned by centrifugation at 500 *g* for 10 min at 4ºC followed by 4000 *g* for 15 min at 4ºC, and filtered with a 0.45 µm filter. The media then were concentrated with a 3kDa cutoff ultrafilter (Millipore, UFC900324), and the protein concentration was determined with Bradford assay (Bio-Rad, Cat # 5000006). Cells were lysed in PierceTM RIPA buffer (Thermo, Cat # 89900) with a protease inhibitor cocktail (Thermo) and the protein concentration was tested with Bradford assay. 75 µg of each sample was provided to the Mass Spectrometry-Based Proteomics Resource Facility at Department of Pathology, University of Michigan for TMT labeling, LC-MS/MS and bioinformatics analysis.

For TMT labeling, cell and medium samples were proteolysed and labeled with TMT 10-plex essentially by following manufacturer’s protocol (ThermoFisher, Cat # 90110, Lot # VJ306782). For liquid chromatography-mass spectrometry (LC-MS) analysis, as previously described (Tank et al., 2018). An Orbitrap Fusion (Thermo Fisher Scientific) and RSLC Ultimate 3000 nano-UPLC (Dionex) was used to acquire the data. 2 µl of the sample was resolved on a PepMap RSLC C18 column (75 µm i.d. x 50 cm; Thermo Scientific) at the flowrate of 300 nl/min using 0.1% formic acid/acetonitrile gradient system (2-22% acetonitrile in 150 min;22-32% acetonitrile in 40 min; 20 min wash at 90% followed by 50 min re-equilibration) and directly spray onto the mass spectrometer using EasySpray source (Thermo Fisher Scientific). Mass spectrometer was set to collect one MS1 scan (Orbitrap; 120K resolution; AGC target 2×105; max IT 100 ms) followed by data-dependent, “Top Speed” (3 seconds) MS2 scans (collision induced dissociation; ion trap; NCE 35; AGC 5×103; max IT 100 ms). For multinotch-MS3, top 10 precursors from each MS2 were fragmented by HCD followed by Orbitrap analysis (NCE 55; 60K resolution; AGC 5×104; max IT 120 ms, 100-500 m/z scan range).

Data analysis was performed using Proteome Discoverer (v2.4; Thermo Fisher). In data analysis, MS2 spectra were searched against SwissProt human protein database (20353 entries; downloaded on 06/20/2019) with the search parameters: MS1 and MS2 tolerance were set to 10 ppm and 0.6 Da, respectively; carbamidomethylation of cysteines (57.02146 Da) and TMT labeling of lysine and N-termini of peptides (229.16293 Da) were considered static modifications; oxidation of methionine (15.9949 Da) and deamidation of asparagine and glutamine (0.98401 Da) were considered variable. Identified proteins and peptides were filtered to retain only those that passed ≤1% FDR threshold. Quantitation was performed using high-quality MS3 spectra (Average signal-to-noise ratio of 10 and <50% isolation interference).

### Quantification and statistics

In all figures, the quantification results are expressed as the mean ± SEM (standard error of the mean) from three independent experiments, unless otherwise stated. The statistical significance of the results was assessed using a two-tailed Student’s *t*-test. *, p<0.05, **, p<0.01, ***, p<0.001.

## Supporting information

Supplemental Table 1-Secretome

Supplemental Table 2. GO term analysis

## Figure assembly

All figures were assembled using Adobe Illustrator; Figure 6L was drawn with Biorender.

## Supplemental Information

Supplemental Information includes 2 supplemental figures and 2 supplemental tables. Figure S1 demonstrates that GRASP55 is required for the secretion of a subset of neurotoxic proteins in addition to Htt. Figure S2 provides evidence that GRASP55 facilitates Htt secretion by stabilizing p23. Table S1 lists the proteins whose secretion is altered by 55KO in the secretome study. Table S2 lists the selected GO term analysis for the secretome of WT and 55KO cells.

## Acknowledgments

We thank Dr. Masaaki Komatsu and Dr. Lois Weisman for Atg7 KO MEF cell lines. We thank Dr. Venkatesha Basrur and the Mass Spectrometry-Based Proteomics Resource Facility in the Department of Pathology at the University of Michigan for their contribution to the Proteomics analysis. We thank the current and former members of the Wang lab for stimulating discussions and technical support. We also thank Dr. Prabhodh Abbineni and Vi Tang in the Ginsburg lab for their valuable input in the secretome experiment. This work was supported by the National Institutes of Health (Grant R35GM130331) and the Fast Forward Protein Folding Disease Initiative of the University of Michigan to Y. Wang, and a University of Michigan Rackham Predoctoral Fellowship to E. Ahat.

## Author Contributions

E.A. and Y.W. designed the experiments; E.A. performed most of the experiments. S.B. performed the western blot experiments for p23 level analysis and provided technical support for secretome data visualization. J.Z. performed the co-immunoprecipitation experiment to determine p23-GRASP55 interaction. F.V and A.N performed the bioinformatics analysis of the secretome data. W.R performed the signal peptide and secretomeP data search of the secretome. L.S and H.L.P. provided the control and HD mouse brain lysates. E.A. and Y.W. analyzed the data. E.A. and Y.W. made the figures and wrote the first draft; all authors discussed the results and contributed to the final manuscript.

## Supplemental Figures and Figure Legend

**Figure S1.**
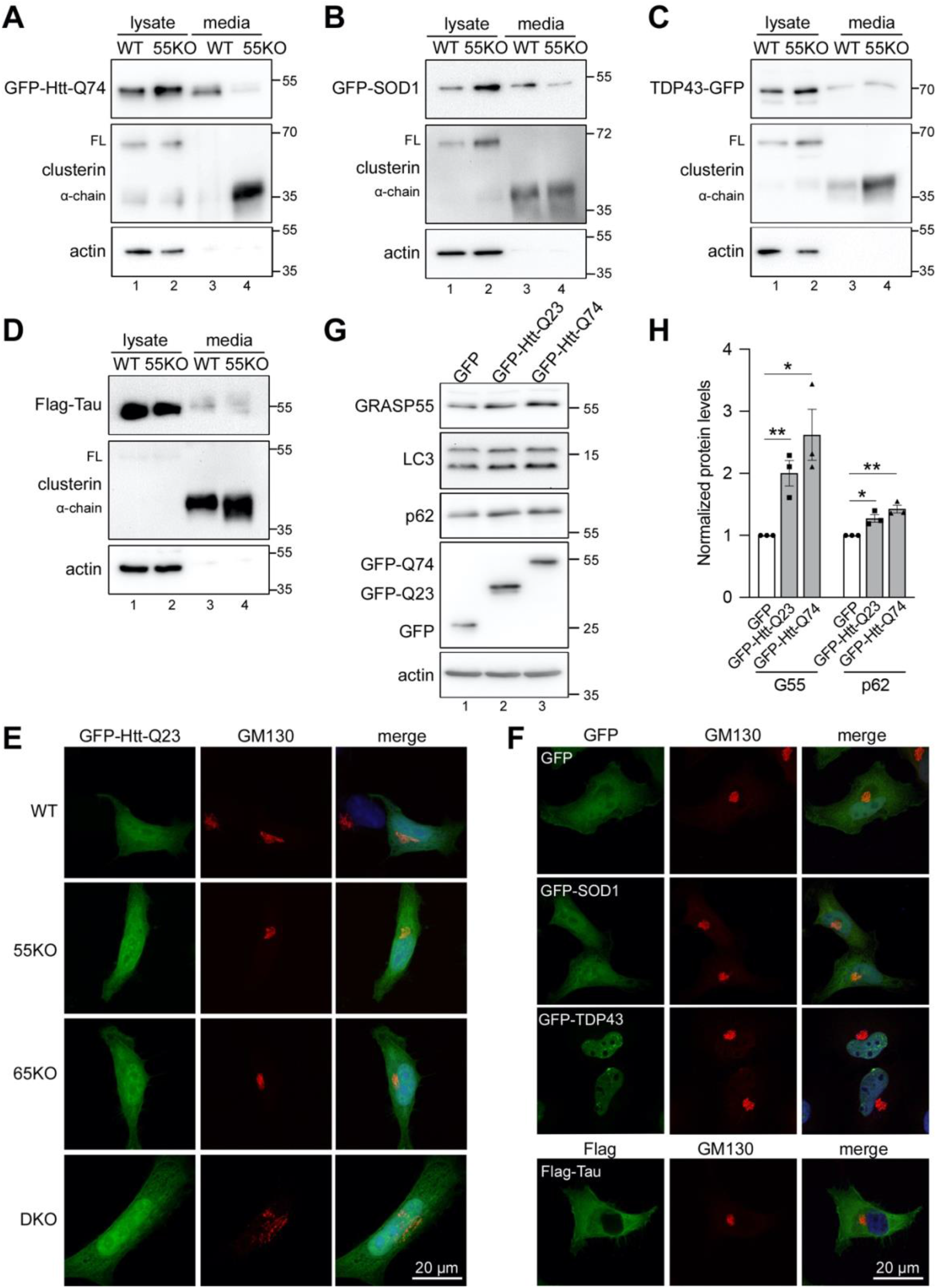
GRASP55 is required for the secretion of a subset of neurotoxic proteins. **(A-B)** Htt and SOD1 secretion is reduced by 55KO. WT and 55KO cells were transfected with GFP-Htt-Q74 (A) or GFP-SOD1 (B) and analyzed for secretion by western blot. Note that 55KO reduced the secretion of both Htt and SOD1. **(C-D)** TDP43 and Tau secretion is GRASP55 independent. WT and 55KO cells were transfected with TDP43-GFP (C) or Flag-Tau (D) followed by the analysis of TDP43 and Tau in the cell lysate and conditioned media. **(E)** GFP-Htt-Q23 expression does not disrupt the Golgi structure. GFP-Htt-Q23 was transfected to WT and GRASP KO cells and stained for GM130. The GM130 signal in DKO cells was increased to better visualize the Golgi morphology. **(F)** Expression of SOD1, TDP43 or Tau does not impact the Golgi structure. WT cells were transfected with indicated constructs and stained for GM130. **(G)** GRASP55 level is upregulated by GFP-Htt-Q74 expression. WT cells expressing GFP, GFP-Htt-Q23 or GFP-Htt-Q74 were analyzed by western blot for indicated proteins. (**H**) Quantification of GRASP55 and p62 levels in G. Results are presented as mean ± SEM; statistical analysis was performed using Student’s *t*-test. *, p < 0.05; **, p < 0.01.

**Figure S2.**
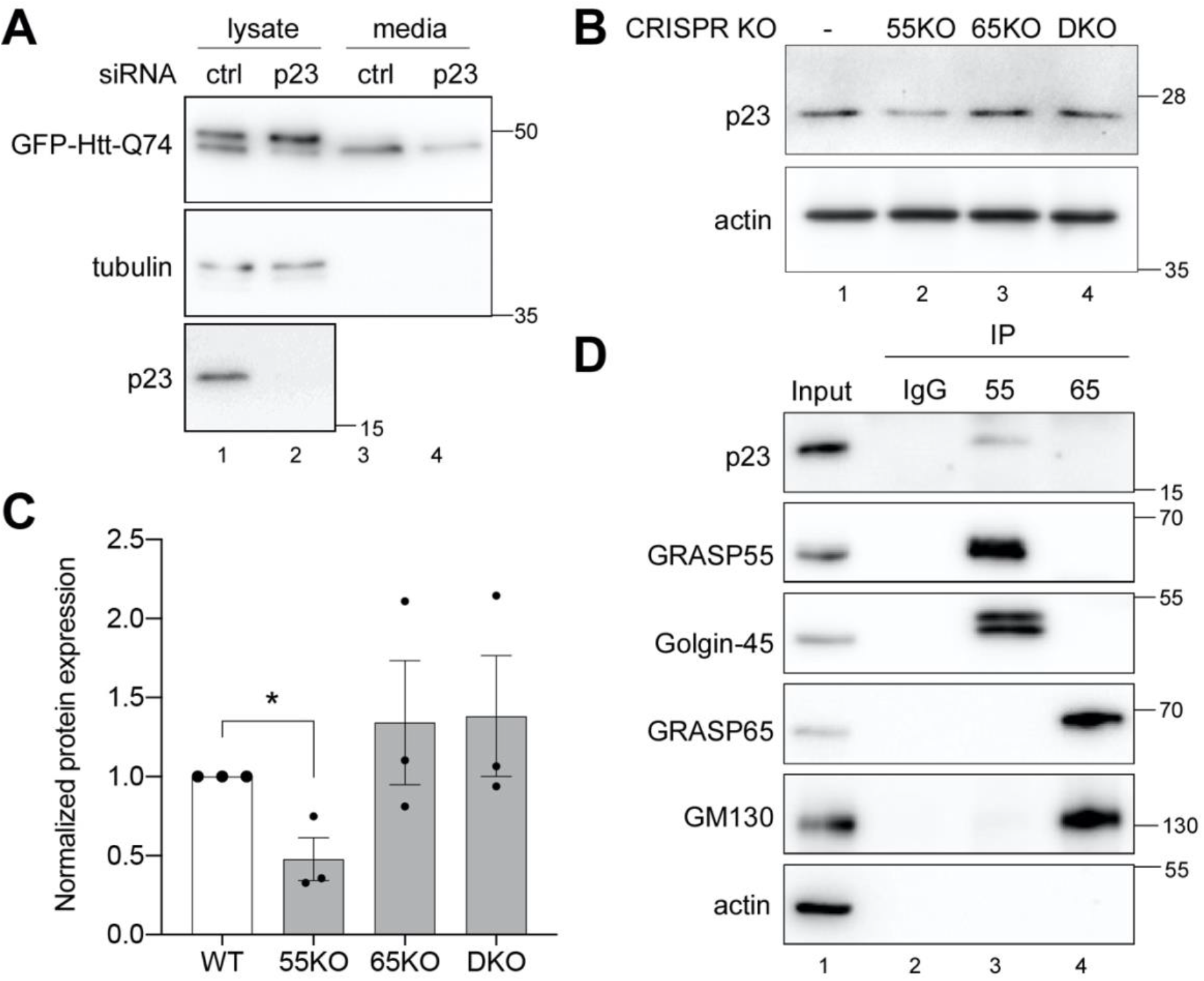
GRASP55 facilitates Htt secretion by stabilizing p23. **(A)** p23 KD reduces Htt secretion. WT HeLa cells were first transfected with control (ctrl) or p23 siRNA for 36 h and then transfected with GFP-Htt-Q74 for 24 h. Htt secretion was analyzed by western blot. **(B)** p23 level is reduced in 55KO cells. Western blot analysis of p23 in WT and GRASP KO cells. **(C)** Quantitation of p23 in B. Results are presented as mean ± SEM; statistical analysis was performed using Student’s *t*-test. *, p<0.05. **(D)** p23 interacts with GRASP55 but not GRASP65. Endogenous GRASP55 or GRASP65 was immunoprecipitated and their interaction with p23 was analyzed by western blot. Golgin-45 and GM130 were used as positive controls for GRASP55 and GRASP65, respectively.

## Supplemental Table

**Table S1. List of differentially secreted proteins in 55KO vs. WT**.

Gene names, fold changes [log2FC (55KO/WT media)], p value, and ER signal sequence are shown. Genes with significant change in 55KO compared to WT (p < 0.05) are manually selected and indicated whether they contain ER signal sequences.

**Table S2. List of selected GO term analysis of WT and 55KO secretome**.

Secreted proteins with significant change in 55KO compared to WT ([|Log2FC (55KO/WT)|>0.5]; p < 0.05) are selected. Protein with or without ER signal sequence are separately processed in Metascape (https://metascape.org/gp/index.html#/main/step1) using the default setting. Top20 significant and relevant GO terms are selected out of Top100 GO terms.

